# Replicated Functional Evolution in Cichlid Adaptive Radiations

**DOI:** 10.1101/2023.09.30.559334

**Authors:** Christopher M. Martinez, Katherine A. Corn, Sarah Williamson, Darien Satterfield, Alexus S. Roberts-Hugghis, Anthony Barley, Samuel R. Borstein, Matthew D. McGee, Peter C. Wainwright

## Abstract

Adaptive radiations highlight the mechanisms by which species and traits diversify and the extent to which these patterns are predictable. We used 1,110 high-speed videos of suction feeding to study functional and morphological diversification in 300 cichlid species from three African Great Lake radiations of varying ages (Tanganyika, Malawi and Victoria) and an older, spatially dispersed continental radiation in the Neotropics. Among African radiations, standing diversity was reflective of time. Morphological and functional variance in Lake Victoria, the youngest radiation, was a subset of that within Lake Malawi, which itself was nested within the older Tanganyikan radiation. However, functional diversity in Neotropical cichlids was often lower than in Lake Tanganyika, despite being at least 25 My older. These two radiations broadly overlapped, but each diversified into novel trait spaces not found in the youngest lake radiations. Evolutionary rates across radiations were inversely related to age, suggesting, at least for lake radiations, extremely rapid trait evolution at early stages. Despite this support for early bursts, other patterns of trait diversity were inconsistent with expectations of adaptive radiations. This work suggests that cichlid functional evolution has played out in strikingly similar fashion in different radiations, with contingencies eventually resulting in lineage-specific novelties.

## Introduction

Adaptive radiations provide glimpses into how traits diversify and evolve across related taxa in the presence of ecological opportunity (Simpson 1953; Gillespie et al. 2020). Studies on adaptive radiations have helped to explain how Caribbean anoles (Losos et al. 1998) and Hawaiian spiders (Gillespie 2004) have colonized new island habitats through repeated evolution of convergent ecomorphs. They have also highlighted mechanisms underlying trait divergence across adaptive peaks in Bahamian pupfishes (Patton et al. 2022), Galapagos finches, and Hawaiian honeycreepers (Tokita et al. 2016). A common theme in the literature on adaptive radiations is the degree to which trait evolution reflects predictable patterns of diversification versus the generation of novel combinations of phenotypes (e.g., Schluter 1996; Losos et al. 1998; Gillespie 2013).

Cichlid fishes are renowned for having multiple expansive radiations involving hundreds of species in each of three large African lakes – Tanganyika, Malawi and Victoria – (Freyer and Illes 1972) and a continental radiation in tropical South and Central America (López-Fernández et al. 2013; Arbour and López-Fernández 2014). The existence of these large radiations of related species provides an opportunity to capitalize on natural replication to address questions about the repeatability of these systems at a scale beyond the first few niche expansions. The radiations also differ considerably in age, approximately 55 Ma for Neotropical cichlids, 28 Ma for Lake Tanganyika (although a recent estimate suggests 10 Ma; Ronco et al. 2021), 2 Ma for Lake Malawi, and 0.1 Ma for Lake Victoria (we discuss cichlid ages further in the supplement). These differences present temporally spaced sample points that allow insight into the long-term unfolding of adaptive radiations and the relative importance of time and rate of diversification on current patterns of diversity.

Previous comparisons of phenotypic diversity in the three African lakes have drawn two main conclusions. First, diversity of body shape and trophic morphology differs between the lakes, with the oldest radiation, in Lake Tanganyika, housing greater diversity of body shape and craniofacial morphology, and the youngest radiation, in Lake Victoria, having the lowest diversity (Young et al. 2009; Cooper et al. 2010). It is not known whether adaptive radiations of the African lakes have amassed greater morphological diversity, and associated functional variation, than the much older continental radiation in the Neotropics that includes roughly 500 species (Lopez-Fernandez et al. 2013). Such a contrast would provide insight into whether the processes of adaptive evolution in the African lakes produces even greater diversity than a continental radiation evolving over a much longer time span. Second, convergent ecology and morphology are common, suggesting relatively predictable modes of diversification and broadly repeated patterns of evolution in feeding morphology (Cooper et al. 2010) and body shape (Kocher et al. 1993; Young et al. 2009). Similar instances of convergence have been found in related groups of Neotropical cichlids (Burress et al. 2017). Thus, large cichlid adaptive radiations could generate similar, though not identical, sets of phenotypes where diversity accumulates over extended time periods.

The temporal sampling of cichlid radiations creates an opportunity to evaluate, at various stages of development, patterns of functional diversification relative to our expectations of adaptive radiations. A classic prediction is an early burst in trait diversification, where evolutionary rates are initially rapid as open niches in newly colonized habitats are filled, followed by a nonlinear decay in rates after initial expansion (Simpson 1953). An assumption of the early burst model is that the rapid increase in trait diversity is achieved by different lineages evolving toward separate adaptive zones where they subsequently undergo further diversification, resulting in comparatively greater variance among clades than within (Simpson 1953; Harmon et al. 2003).

In the present study, we describe the diversity of cichlid prey capture kinematics using high-speed video recordings of 300 species, sampled from the three African great lake radiations and the Neotropics. We contrast kinematic and morphological variation of the feeding mechanism in radiations of varying age to test the hypothesis that differences in standing functional diversity are due to time, as opposed to different rates of evolution. To assess the repeatability and predictability of cichlid adaptive radiation, we also quantify the extent to which each has produced species with similar feeding kinematics. Additionally, we test the key expectation that adaptive radiations exhibit an early burst in trait diversification that is achieved through the partitioning of traits among clades (Simpson 1953; Harmon et al. 2003).

A secondary objective of this work is to examine the relationship between morphological and functional diversity of the cichlid feeding mechanism. Preliminary estimates of body and craniofacial variation in the three large African lake radiations have typically been interpreted as reflecting functional diversity linked to locomotor and feeding biomechanics (Young et al. 2009; Cooper et al. 2010). In many cases, links between morphological variation and functional properties are well-established (Hulsey & García de León 2005; Hulsey et al. 2006; Higham et al. 2007). Nevertheless, our current understanding of functional diversity in cichlids is largely inferred from morphological variation, rather than direct measurements of functional traits. Comparisons of functional diversity allow us to test the reliability of morphological variation to reflect function and help to identify key axes of diversification that are cryptic when only morphology is considered.

## Methods

### Species sampling and feeding videos

1,110 high-speed videos of feeding motions from 300 species of lab-filmed cichlids were studied. Species were broadly distributed phylogenetically from one of four focal radiations, the Neotropics (n=85 species) and African Great Lakes, Tanganyika (n=89), Malawi (n=86), and Victoria (n=40). We note that two species, *Harpagochromis sp.* “golden duck” and *Pyxichromis orthostoma*, belonging to the Lake Victoria Region Superflock are endemic to Lake Kyoga, which retains a connection to Lake Victoria via the Victoria Nile River. All videos were filmed from a lateral perspective at 2,000 frames-per-second (McGee et al. 2016) and contained full-effort suction feeding strikes on moderately evasive, living midwater prey. Primary prey included mosquito larvae (*Culex pipiens*), black worms (*Lumbriculus* sp.), and *Daphnia magna*. Small fish were rarely used as prey to elicit sufficiently full effort feeding strikes from some species, which was important for reducing kinematic variation due to fish effort. We extracted 10 frames from each video, equally spaced in time from the initiation of the motion to peak expansion of the feeding apparatus, prior to mouth closing. For comparative analyses, we matched filmed species to a recent cichlid phylogeny (McGee et al. 2020; Supplementary Information).

### Morphological and functional traits

Cichlid functional diversity was determined using a variety of kinematic traits, all derived from an initial configuration of 18 cranial landmarks manually placed on each of 10 video frames for a motion (fig. 1A). Digitizing was done in tpsDIG2 (Rohlf 2015) and StereoMorph (Olsen and Westneat 2015). Landmark data for 53 species came from a previous study (Martinez et al. 2018) but the remaining 247 species comprised new data. First, subsets of landmarks were used to measure movements (i.e., maximum excursions) of key morphological features involved in prey capture. In total, we created six *motion component* traits (fig. 1B), three from rotational movements of bones (*lower jaw rotation*, *cranial rotation*, *maxillary rotation*) and three from linear displacements (*premaxilla protrusion*, *hyoid depression*, *mouth gape*). We analyzed traits both as multivariate *motion components*, and individually. Due to the incommensurability of angles and linear displacements (Huttegger and Mitteroecker 2011), we converted the three rotational traits to distances by using the observed angle of rotation and the length of the rotating arm (measured on the fish at full gape) to determine the length of the arc transcribed by the structure in question (e.g., the Euclidean distance travelled by the distal end of the maxilla). All component traits were then scaled by dividing values by the centroid size of the fish’s head in a closed-mouth state. Lastly, we averaged the components across repeated feeding trials within individuals, and then across individuals to get a mean trait value for the species. All additional traits described below were similarly averaged to species for comparative analyses.

**Figure 1.**
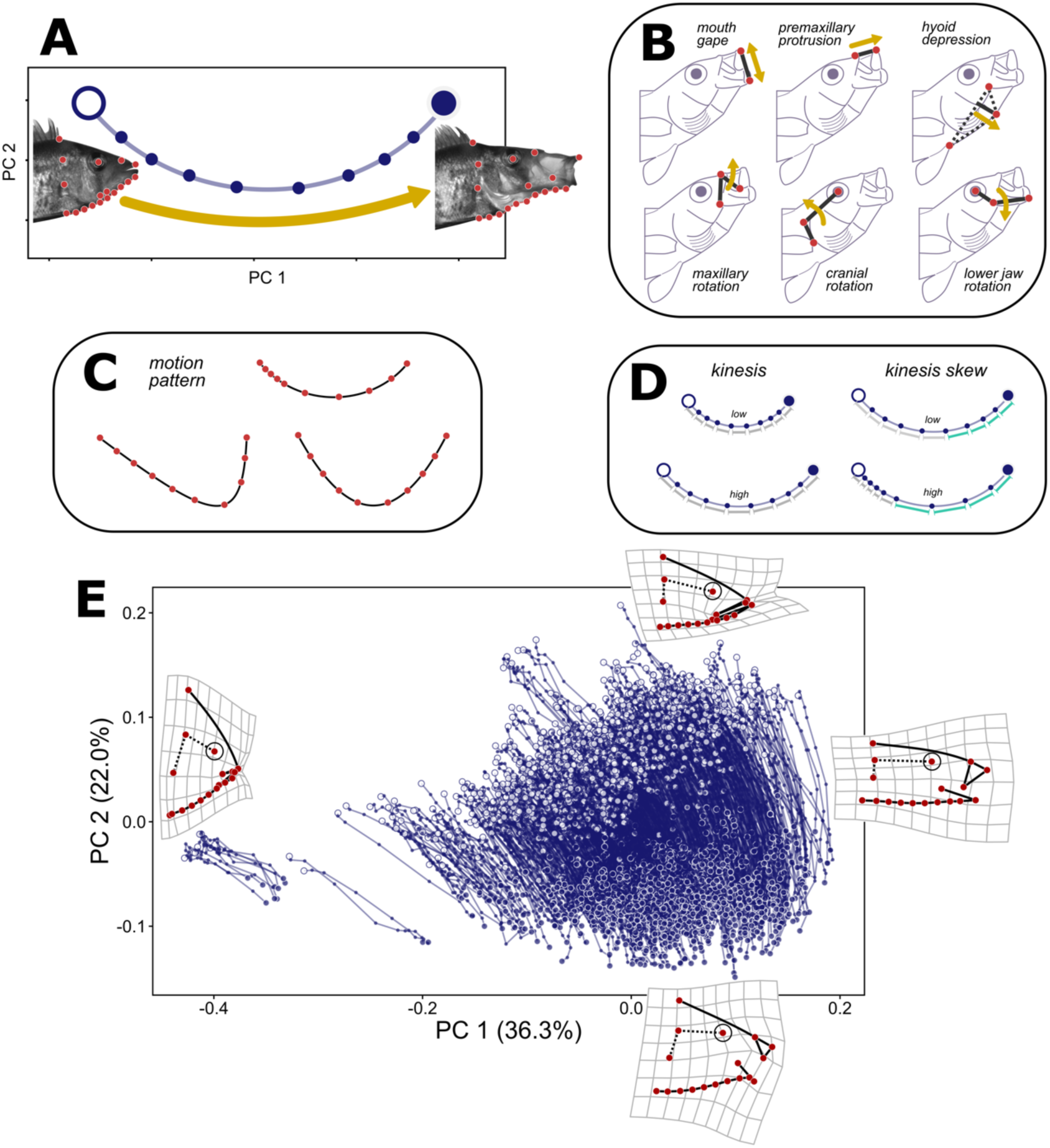
Functional traits examined across 300 species of cichlid in this study. **A**) Principal component axes, PC 1 and PC 2, displaying shape change for a single trajectory of 10 cranial shapes during a suction feeding motion. **B**) Subsets of two or three landmarks were used to measure maximum excursions of six commonly measured *motion components* of suction feeding in percomorph fishes. **C**) Changes in the timing or extent of movements represent differences in *motion pattern* that are manifested as variation in trajectory shapes. Trajectories varied in spacing of cranial shape changes and the relative symmetry of trajectory paths. **D**) A series of trajectory-derived traits related to mobility, included cranial *kinesis* (total trajectory length), *kinesis coefficient* (not pictured, total output kinesis divided by input movement from cranial rotation), and *kinesis skew* (kinesis over the last five motion shapes divided by total kinesis). **E**) PCs 1 and 2 from 1,110 trajectories of suction feeding motions, with deformation grids displaying shape change. All plots are shown in two dimensions for visualization, but data analysis was in the full dimensionality of the shape data unless otherwise noted.

Additional kinematic traits were created using an approach that characterizes movements as trajectories of shape change (fig. 1), integrating the numerous moving parts involved in a complex motion into a single object that allows for comparisons at the whole-motion level (Adams and Cerney 2007; Adams and Collyer 2009; Collyer and Adams 2013; Martinez et al. 2018; 2022; Martinez and Wainwright 2019). Digitized cranial landmarks were aligned and scaled using generalized Procrustes analysis (GPA) with the ‘gpagen’ function in the ‘geomorph’ package, v 4.0.3, in the R statistical environment, v 4.1.3 (Adams et al. 2021; R Core Team 2022), with alignment of sliding semi-landmarks along the ventral margin of the head achieved by minimizing Procrustes distances. Once aligned with GPA, the progressive movements of landmark-tracked cranial features result in a trajectory of shape change (fig. 1A & E), the features of which can be used as traits that capture motion variation (Martinez et al. 2018; 2019; 2022). In this study, for example, the length of each motion trajectory is a measure of cranial *kinesis*, or the amount of movement generated by the feeding apparatus during prey capture (fig. 1D). The total trajectory length was computed as the sum of Procrustes distances between consecutive motion shapes (Collyer and Adams 2013).

We also generated two composite traits designed to provide context about when and how kinesis is achieved. *Kinesis skew* was the natural logarithm of the ratio of kinesis across the final five motion shapes to the total kinesis for the motion. It is a descriptor of the temporal distribution of kinesis within a movement, with smaller values indicating comparatively more movement toward the beginning of the feeding strike and larger values meaning that movement is concentrated near the end of the strike. Next, we measured a *kinesis coefficient* trait, as an analog to kinematic transmission (Westneat 1994; 2004), which is commonly used with biomechanical linkage models to describe output movement of an anatomical feature, given a degree of input motion from another. Here, we took the natural logarithm of total kinesis for a motion (output movement) divided by maximum cranial rotation (input movement) from the *motion components* described above. We used cranial rotation, which is driven by contraction of epaxial muscles posterior to the head, for the input value as it facilitates expansion of the buccal cavity and drive movements of other features of the feeding apparatus (Camp et al. 2020).

The final functional trait we compared was *motion pattern*, briefly described here. For complex biomechanical systems composed of numerous mobile features, any change in relative timing and/or degree of movement across those features causes variation in the pattern of movement at the whole-motion level. We used anatomical landmarks to express feeding movements as an ordered series of changing shapes over time, or a trajectory through morphospace (fig. 1A). The paths forged by these trajectories each have their own shape that reflects motion patterns – different trajectory shapes represent different patterns of movement that can be observed both within a single species feeding with different modes of prey capture (Martinez et al. 2022) and across species with different evolved feeding systems (Martinez et al. 2018). We note that *Motion components* and *motion pattern* are both multivariate descriptors of feeding movements but capture contrasting aspects of their diversity. *Motion components* measure maximum excursions of key features of feeding motions, whereas *motion pattern* describes how and when those movements take place (Martinez et al. 2022). To compare *motion patterns*, we used modified code from the ‘trajectory.analysis’ function in the R package ‘RRPP’ (version 1.0.0) to align and scale trajectories (fig. S1; Collyer and Adams 2018; 2019). Here, the centroid size of the entire trajectory was the scaling factor.

To provide context to functional and kinematic patterns, we also examined interspecific cranial morphologies across cichlid species. We extracted head shape data from the starting positions of motions, where the mouths were in a closed state. A separate shape alignment was done on head shape landmarks, which were then averaged to species prior to statistical analyses.

### Trait diversity and overlap among radiations

Variance of morphological and kinematic traits, both univariate and multivariate, were measured and statistically compared using the ‘morphol.disparity’ function in ‘geomorph’ with 10,000 permutations. In addition to trait variation, we were interested in the degree of overlap (or lack thereof) in the occupation of multivariate functional and morphological spaces across cichlid radiations. We created four-dimensional hypervolumes for *motion components*, *motion pattern*, and head shape data using the R package ‘hypervolume’ v 3.0.0 (Blonder et al. 2014; 2018). We took the first four axes from a principal component analysis, as hypervolumes are best conducted on orthogonal variables (Blonder et al. 2014; 2018). Hypervolumes were made for each radiation (e.g., species from Lake Tanganyika), and for subsets of the data excluding each radiation (e.g., all species *not* from Lake Tanganyika). We then assessed hypervolume overlap and the fraction of unique space occupied by each radiation. Lastly, we estimated the likelihood of our observed results against a null distribution of hypervolumes generated by randomly permuting group assignments among species 10,000 times (e.g., Corn et al. 2022).

### Rates of evolution

For all traits, we estimated rates of evolution (the Brownian rate parameter) within each cichlid radiation with the ‘compare.evol.rates’ function in ‘geomorph’ for both univariate and multivariate rates, with significance based on 10,000 permutations.

### Modes of trait diversification

We examined both historical reconstructions of trait diversification and contemporary patterns of variation to explore the manner by which trait diversity was attained in cichlid radiations. We estimated the accumulation of trait disparity through time (DTT) in radiations using the ‘dtt’ function in the R package ‘geiger’ v 2.0.7. (Harmon et al. 2008; Pennell et al. 2014). DTT plots show relative disparities among subclades at each divergence event in the tree, estimating whether trait diversity is concentrated within or among subclades as an explicit test of the early burst expectation (Simpson 1953). An output of this analysis is the morphological disparity index (MDI), a metric for comparing the difference between the estimated relative disparity of a clade and the disparity of the clade under simulated Brownian motion. MDI statistics were calculated for the first 75% of the tree’s history, as missing species in the recent phylogeny may obscure patterns close to the present. We estimated DTT from subtrees of each radiation for *motion components*, *motion pattern* and head shape, using the first four axes from PCAs on each, consistent with our comparisons of hypervolumes. Tree topology is key to DTT analyses, so we note caution in interpreting results for young radiations, like Lakes Malawi and Victoria, in which a tree-like model of lineage diversification is unlikely due to widespread hybridization (Joyce et al. 2011; Meier et al. 2017; Scherz et al. 2022). Consequently, we focus our discussion on the two older radiations in Lake Tanganyika and the Neotropics but provide results for all radiations in the supplement materials.

To further examine patterns of trait dispersion, we computed distances between extant species and radiation-specific ancestral states. For each cichlid radiation, we estimated ancestral states under Brownian motion with the ‘gm.prcomp’ function in ‘geomorph’, extracting the value at the root of the tree as the most recent common ancestor (MRCA) for the radiation. Finally, we measured Euclidean distances (*motion components*) and Procrustes distances (*motion pattern* and head shape) between each species and its radiation’s MRCA.

## Results

### Functional diversity across cichlid radiations

*Motion components*, composed of six key features of fish cranial movement during feeding (fig. 1B), displayed 3.2 (Lake Tanganyika) and 2.9 times (Neotropics) greater variance in older radiations compared to the youngest radiation in Victoria (figs. 2 & 3; table S3). Separate univariate analyses on the individual components did show some variation in rank orders of variances across traits (fig. 2; table S3). In all cases, Lake Victoria had the lowest variance, followed closely by Lake Malawi, but some traits displayed their highest variation in Lake Tanganyika (*premaxillary protrusion*, *maxillary rotation*, *lower jaw rotation*, and *mouth gape*), while others were most variable in the Neotropics (*cranial rotation* and *hyoid depression*).

**Figure 2.**
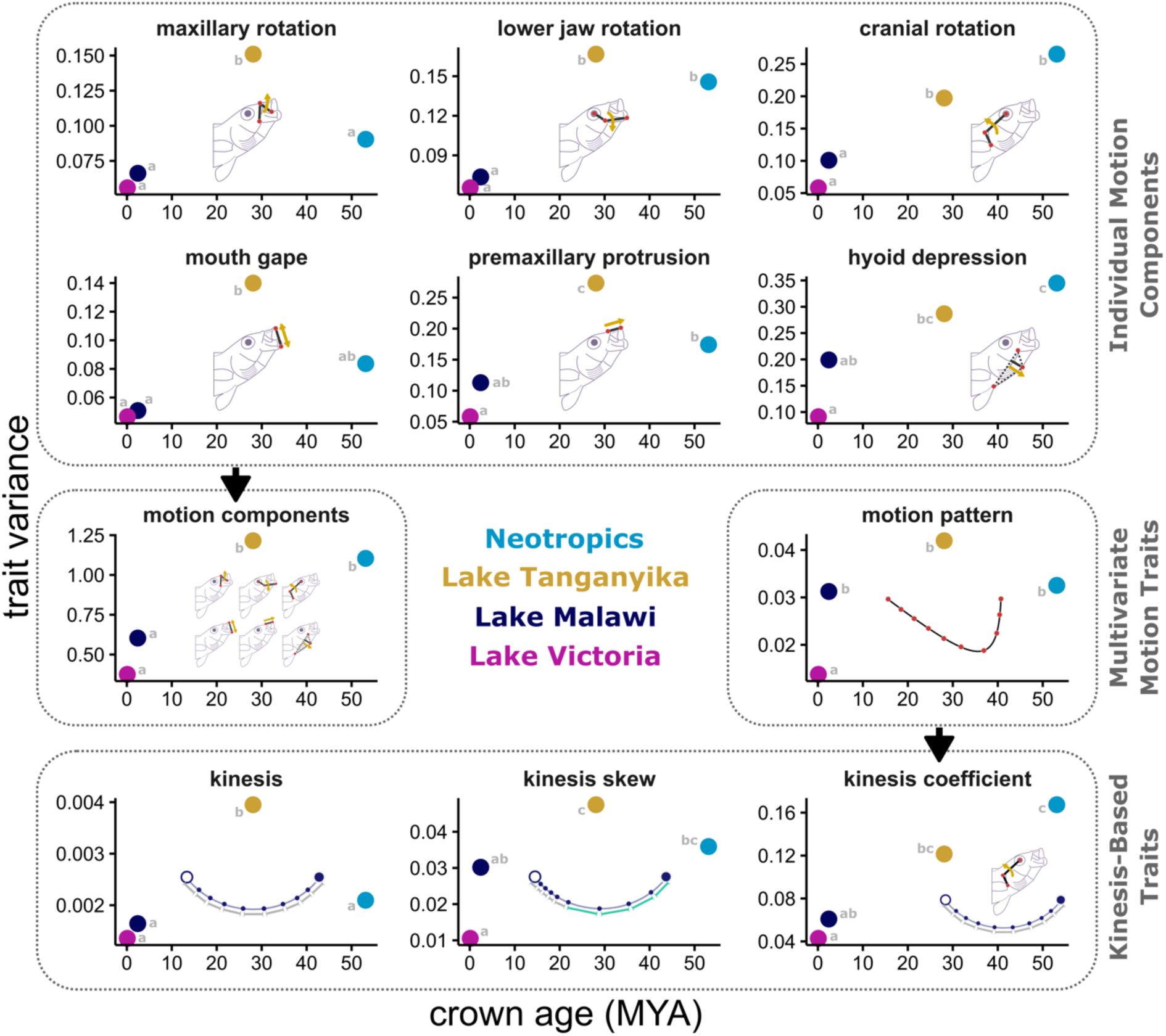
Functional trait variances in cichlid adaptive radiations plotted against crown age. Six individual kinematic traits are shown on top, with an arrow pointing to the multivariate *motion components* containing all of them. *Motion pattern* (i.e., the shape of a kinematic trajectory) is shown in the middle with an arrow pointing to composite *kinesis-based* traits measured from trajectories. Letters next to plot points denote significant P-values from pairwise comparisons. Radiations sharing a letter do not have statistically different variances.

**Figure 3.**
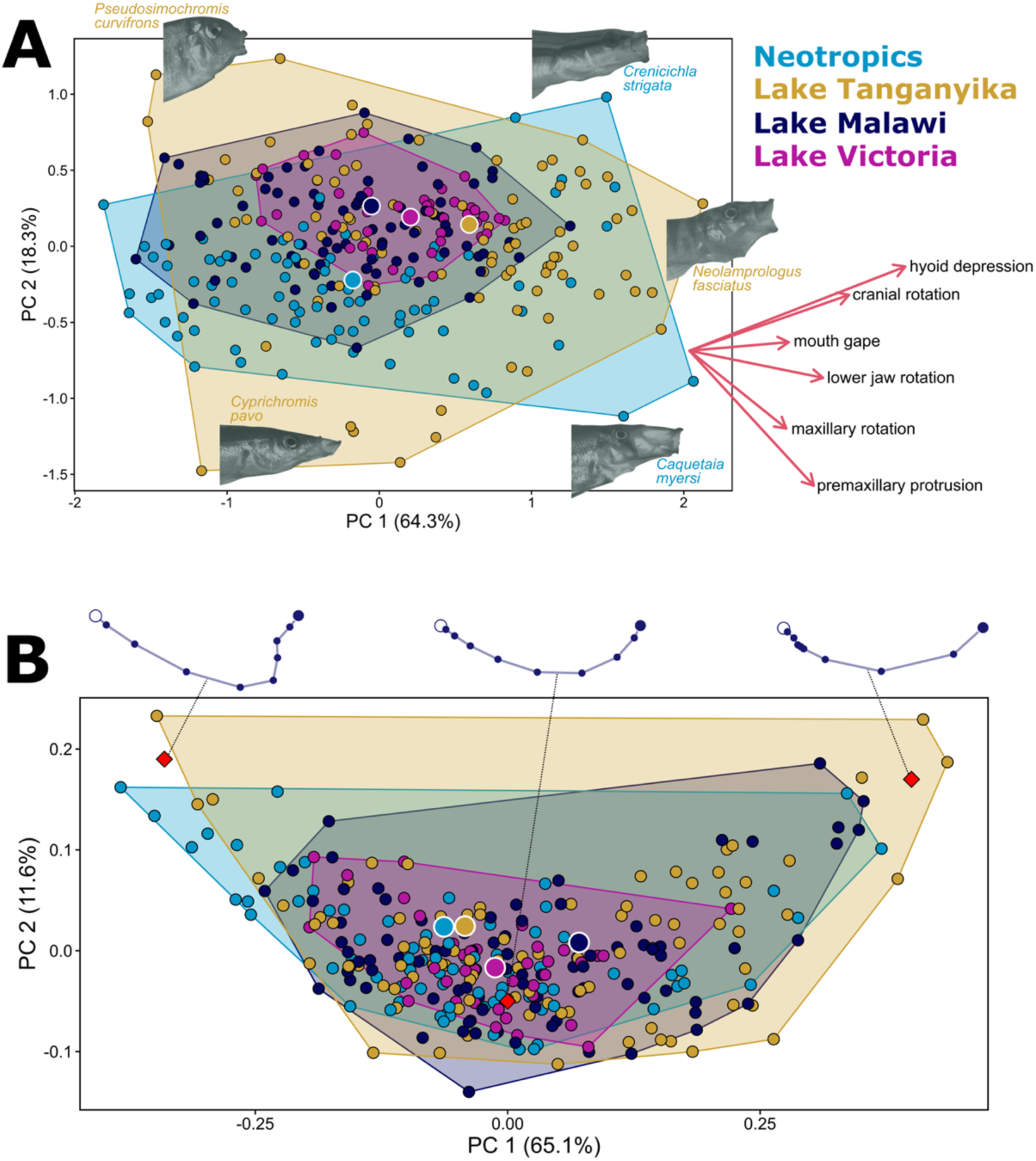
Primary dimensions of variation, PCs 1 and 2, from separate principal component analyses for species-averaged multivariate functional traits. **A**) *Motion components*, a dataset comprising measurements of maximum excursions for six key kinematic features are plotted by cichlid radiation. Cichlid heads at maximum gape are shown across the plot, and trait loadings are shown on the right. **B**) Variation in *motion patterns* by radiation, with two-dimensional representations of the shapes of kinematic trajectories provided at select locations of the plot (red diamonds). Low values on PC 1 are motions with an abrupt shift in the manner of cranial movements and large values are motions in which kinesis is disproportionately concentrated toward the end of the strike. For both multivariate traits, Lake Tanganyika and the Neotropics display unique occupation of this space, while Lakes Malawi and Victoria are almost entirely nested within them. Estimated values of most recent common ancestors (MRCA) are shown as large dots with white borders.

Functional traits derived from trajectories of shape change also showed different levels of diversity across radiations. Cranial *kinesis* had significantly greater variance in Lake Tanganyika than all other cichlid radiations, having 1.9-times greater variance than the next closest (the Neotropics) and 2.9-times greater variance than the lowest (Lake Victoria). *Kinesis skew*, the proportion of kinesis in the latter half of the feeding motion, was again most variable in Lake Tanganyika, but there was greater parity between it, Lake Malawi and the Neotropics (fig. 2). *Kinematic coefficient*, output kinesis relative to input movement from cranial rotation, showed the highest variance in the Neotropical radiation, which was significantly greater than that of the younger radiations in Lakes Malawi and Victoria (fig. 2). Finally, the rank order of variances for whole *motion pattern*, a trait describing the timing and sequence of motion events, was the same as they were for *kinesis skew* – Lake Victoria was significantly lower than the other major radiations, followed by nearly identical values in Lake Malawi and the Neotropics, and the greatest variance again in Lake Tanganyika (fig. 2 & 3). Despite its young age, Lake Malawi showed surprisingly high diversity in *motion pattern*. This appears to be driven by a collection of species like *Tyrannochromis nigriventer*, *Copadichromis virginalis*, and *Caprichromis liemi* that have feeding motions in which kinesis is disproportionately concentrated toward the end of the strike (i.e., high *kinesis skew*, and *motion patterns* with large positive PC 1 scores in fig. 3B).

### Occupation of novel functional spaces

Comparisons of hypervolumes identified whether individual radiations occupied unique regions of multivariate functional spaces. For the *motion components*, we found that 62% of Lake Tanganyika’s and 49% of the Neotropics’ functional space was unique to those radiations at the exclusion of all others, with both results occurring in the 99^th^ percentile of randomized trials (fig. 3A; table S4). Cichlids occupying unique regions of the *kinematic components* functional space in Tanganyika included several specialized planktivores with high upper jaw protrusion in the genus *Cyprichromis* and elongate species with comparatively small gapes, like *Chalinochromis brichardi* and *Julidochromis dickfeldi*. In the Neotropics, multiple species of *Amatitlania* and a host of species in the tribe Geophagini (all with relatively small mouths, low cranial rotation, and low hyoid depression) occurred in one unique region, and species capable of extreme upper jaw protrusion, *Petenia splendida* and *Caquetaia myersi*, were found in another. In contrast to Tanganyika and the Neotropics, only 6% of this space in Lake Malawi and 3% in Lake Victoria was unique to those lakes, suggesting that most of their diversity is nested within the other cichlid radiations (fig. 3A; table S4).

Across all species, the diversity of *motion patterns* was largely restricted to a distinct concave or arched distribution (fig. 3B). This reflects a general constraint on motion diversity – despite differences in the relative magnitude of movements (*motion components*), patterns of feeding movements in cichlids are created by the same morphological features, moving in the same direction, and mostly in the same sequence. One extreme of the *motion pattern* distribution was characterized by an abrupt shift in the direction of cranial shape change within morphospace toward the end of the motion, which manifested as an asymmetrical trajectory shape (left side of fig. 3B). During feeding sequences, this was caused by late onset cranial rotation and hyoid depression after full protrusion of the upper jaw was achieved, presumably for continued buccal expansion posteriorly to prolong suction during prey acquisition. The other extreme contained symmetrically shaped trajectories in which the rate of cranial shape change was slow initially but fast toward the end of the strike, such that kinesis was disproportionately concentrated in later motion stages (right side of fig. 3B), a pattern reminiscent of high *kinesis skew*. Comparisons of hypervolumes for *motion pattern* revealed a mostly nested pattern in which Tanganyika (54%) occupied the greatest volume of unique space (upper 99^th^ percentile of permutations), with much lower values for the Neotropics (26%), Malawi (14%), and Victoria (1%; table S4).

### Rates of functional evolution

Across all functional traits, there was a strong inverse and nonlinear relationship between rates and crown ages of radiations (fig. S2; table S5). In each case, Lake Victoria had much higher rates than other radiations, ranging anywhere from 42 to 95-fold faster diversification compared to the radiation with the slowest rate. Similarly, the second youngest radiation, Lake Malawi, consistently had the second highest rates of trait evolution. Pairwise comparisons of rates between Lakes Victoria and Malawi were statistically significant at the α=0.05 level for functional traits except for *hyoid depression* and *kinesis skew*. One caveat is that these analyses assume a tree-like pattern of lineage diversification, and so we limit our interpretation primarily to emphasize the vastly different time scales over which functional trait diversity has accumulated in these radiations. In comparisons between the two older radiations, rates in Lake Tanganyika were always higher than in the Neotropics, ranging from 1.1-fold to 4-fold differences. However, unlike other pairwise comparisons among radiations, Tanganyika and the Neotropics failed to show significant differences with each other for over half of the functional traits considered. Significant differences in rate of evolution in these two radiations were found for *premaxillary protrusion*, *kinesis skew*, *kinesis*, *motion components*, and *motion pattern*.

### Modes of functional diversification

Disparity through time (DTT) analyses provided information about temporal patterns of diversification for two multivariate functional traits, *motion components* and *motion pattern*. In Lake Tanganyika and the Neotropics, neither trait displayed DTT trends that were statistically different from Brownian motion – the morphological disparity index (MDI), describing the deviation of observed DTT trends from the null expectation, was not statistically significant for either dataset (fig. 4A; see table S6 for results from all radiations). However, while DTT trends for *motion components* largely stayed within the 95% range of simulated trait histories, those for *motion pattern* were above the Brownian expectation in all radiations, in some cases for prolonged durations, suggesting that trait variance was at times disproportionately concentrated within subclades. Notably, none of the functional DTT trends fell below the lower 95% range for Brownian motion, an indicator of early bursts of trait diversification during adaptive radiation (fig. 4C).

**Figure 4.**
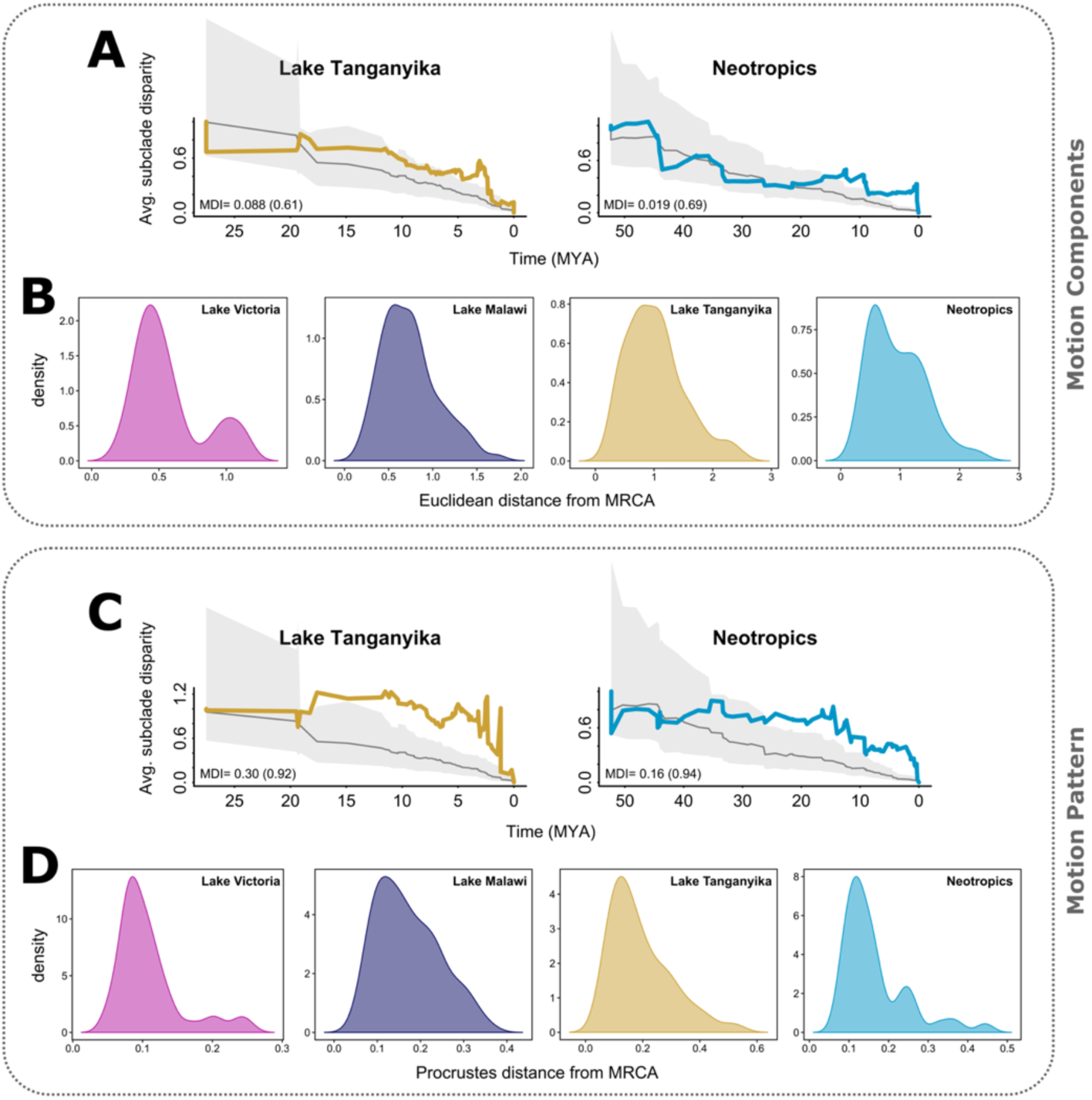
Disparity through time (DTT) plots in the two oldest cichlid radiations, Lake Tanganyika and the Neotropics, for **A)** *motion components* and **C)** *motion pattern*. Also shown are distributions of **B)** Euclidean distances *of motion components* and **D)** Procrustes distances of *motion pattern* between extant species and their radiation-specific most recent common ancestor (MRCA).

Trait dispersion of extant species in each radiation around their most recent common ancestor (MRCA) additionally captured patterns of trait space occupation across radiations at different stages of development. Euclidean distances of *motion components* between species and the MRCA were continuously distributed for all radiations except Lake Victoria (fig. 4B). In this youngest lake, a handful of species with comparably low upper jaw protrusion formed a small secondary peak that was more distantly situated from the Lake Victoria radiation MRCA (observations in the upper left of the Lake Victoria distribution in fig. 3A). These species consisted mostly of herbivorous cichlids from the genus *Neochromis* and omnivores in the genus *Pundamilia*, possibly representing a (weakly) isolated adaptive peak related to trophic ecology and jaw function. For *motion pattern*, Procrustes distances from radiation-specific MRCAs were right-skewed, particularly in Lake Victoria and the Neotropics, seemingly reflecting the highly constrained distribution of the trait more than distinct adaptive zones (figs. 3B & 4D).

### Cranial morphology and its relation to motion diversity

Interspecific variance in head shape was highest in Neotropical cichlids, but only marginally greater than in Lake Tanganyika (table S3). Still, the Neotropics boasted 2.3-times more head shape diversity than Lake Malawi and 2.7-times more than Lake Victoria. Pairwise comparisons of variances were statistically significant except between the two youngest radiations, Malawi-Victoria, and the two oldest, Neotropics-Tanganyika (table S3). Interestingly, the high head shape variance in the Neotropics did not directly translate to functional diversity, as Lake Tanganyika still had greater (but similar) diversity in *motion components*, and significantly higher variance in *kinesis*, *motion pattern, premaxillary protrusion*, and *maxillary rotation* (fig. 5C-E).

**Figure 5.**
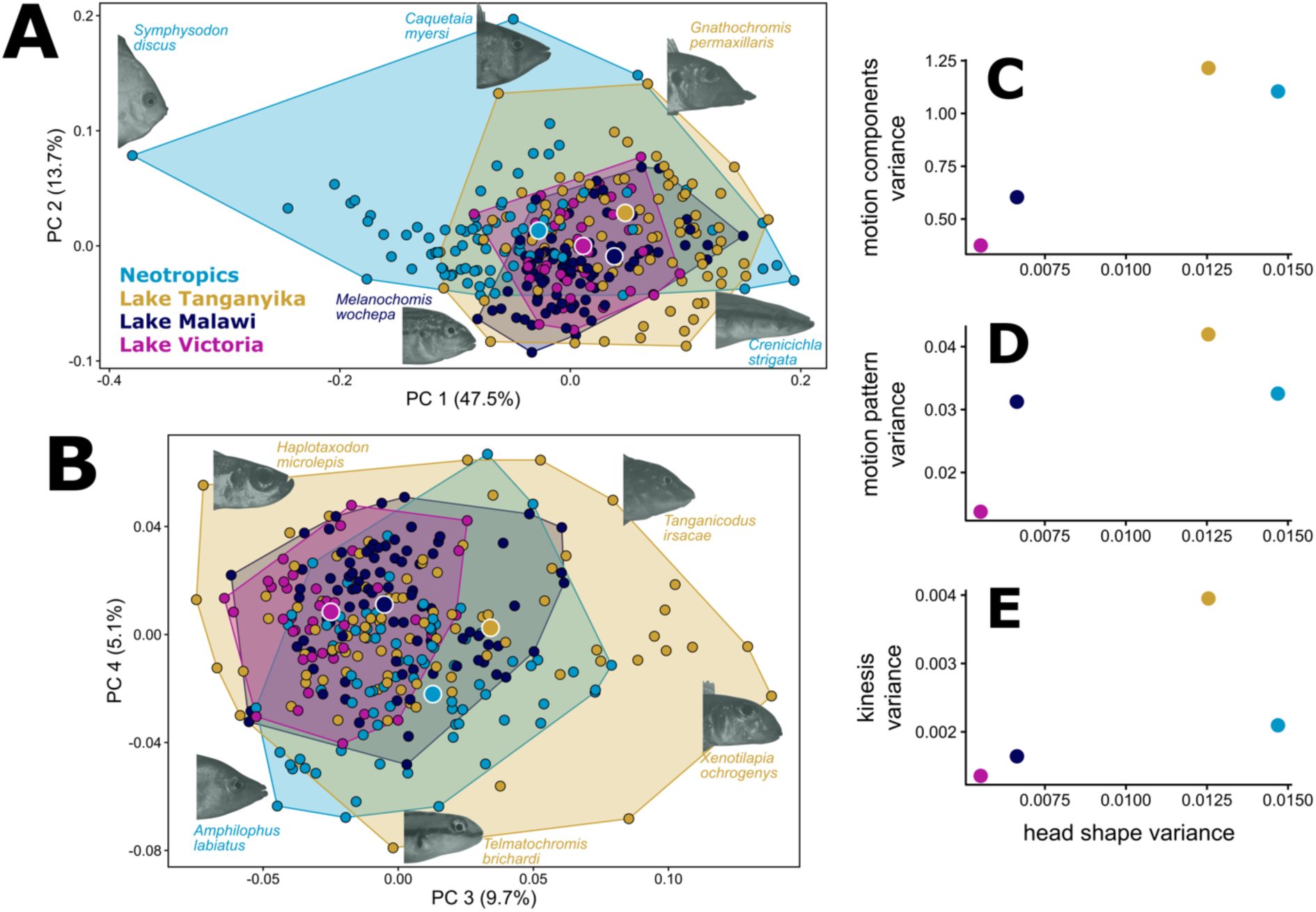
Species-averaged interspecific head shape diversity across cichlid radiations. **A)** Major axes of variation, PCs 1 & 2, show the wide diversity of head shapes in Neotropical cichlids, most notably in head depth (left of plot). Small-mouthed taxa in African lakes, often benthic biting feeders, are concentrated toward the bottom of the plot. **B**) PCs 3 and 4 display axes on which Lake Tanganyika contains high morphological diversity, including variation in orientation of jaws. **C-E**) variance in head shape is shown in relation to select kinematic traits, displaying relationships between form and function. High morphological diversity in the Neotropics does not translate to commensurate kinematic diversity. Most recent common ancestors (MRCA) are shown as large dots with white borders.

Comparisons of hypervolumes for head shape revealed that 59% of cranial diversity in the Neotropics and 57% in Lake Tanganyika was unique to those regions, both observations occurring in the upper 99^th^ percentile of randomized permutations (table S4). One of the things that made the Neotropics stand out was expansion towards deep-headed taxa, like *Symphysodon discus* and *Pterophyllum scalare*, but also several species in the genus *Amatitlania* that were not as extreme but still outside of the space occupied by cichlids in other radiations (fig. 5A, lower scores on PC 1). Some species from the African lake radiations occurred in a non-overlapping region of morphospace with the Neotropics that contained many small-mouthed benthic biting and picking specialists, like *Tropheus brichardi*, *Melanochromis wochepa*, and *Chalinochromis popelini* (fig. 5A, lower scores on PC 2). In addition, Lake Tanganyika possessed a fair degree of unique morphologies varying broadly in direction of mouth orientation (fig. 5B), from upturned (e.g., *Haplotaxodon microlepis*) to downward deflecting profiles (e.g., *Xenotilapia ochrogenys*). Morphological diversity within Lakes Malawi and Victoria was almost entirely contained within the other regions, with only 5% and 3% unique morphospace occupation, respectively.

Rates of head shape evolution were again lowest in the older radiations (Neotropics and Lake Tanganyika), faster in Lake Malawi and much faster in Lake Victoria (fig. S2; table S5). All pairwise comparisons of rates were statistically significant (table S5). Like functional analyses, disparity through time (DTT) trends for head shape were statistically indistinguishable from a Brownian process (fig. S3; table S6). However, in the Neotropics the DTT trend dipped just below the 95% range for Brownian simulations briefly from 44-40 Mya, and again for an extended time from about 39 to 28 Mya (fig. S3). During these periods, head shape disparity was concentrated among clades at a level beyond the null expectation, likely representing evolution of clades towards different adaptive peaks.

## Discussion

In this study, we provide the first quantitative comparison of functional diversity across four major cichlid radiations, leveraging the largest-ever comparative kinematics dataset of vertebrate organisms to contrast patterns of adaptive diversification across vastly different temporal scales (many thousands of years to over 50 My) and spatial ranges (individual lakes versus continental scale). We show that standing functional diversity in African cichlids is strongly related to radiation age and displays a striking nested pattern in which trait spaces occupied in Lakes Victoria and Malawi were almost fully contained within that of Lake Tanganyika. Somewhat surprisingly, functional variance in the much older, continental Neotropical radiation was lower than it was in Lake Tanganyika for many traits, making the high diversity in the latter all the more impressive. This suggests that diversifying forces have operated more effectively in Lake Tanganyika, and likely across the African Great Lakes, as compared to the largely riverine cichlids of Central and South America. Indeed, numerous cichlid lake radiations beyond those examined in this study experience elevated rates of morphological evolution (Burress and Muñoz 2023).

### Cichlid feeding systems and adaptive radiation

Cichlids have long served as a model system for understanding adaptive radiation (Stiassny and Meyer 1999; Seehausen 2006; Turner 2007), yet we recovered mixed evidence that functional diversification of their feeding systems adheres to traditional expectations of this process (Simpson 1953). A negative relationship was found between radiation age and rates of kinematic and morphological evolution (fig. S2), suggesting that phenotypic diversification proceeds fastest in early-stage radiations in a manner consistent with an early burst. Young cichlid radiations in Lakes Victoria and Malawi support modest levels of trait diversity, but they have acquired it at an incredibly fast pace. Although there may be some effect in our study of the natural time-dependency of macroevolutionary rate estimates (Harmon et al. 2021), the relationship between disparity and age of each radiation (fig. 2) strongly suggests a negative correlation between age and rate of phenotypic diversification.

In addition to an early burst, an assumption of adaptive radiations remains that at some stage of early diversification, trait variance will be distributed disproportionately among clades versus within them (Simpson 1953; Harmon et al. 2003). Though cranial morphology showed some hints of elevated divergence among lineages in the Neotropics (fig. S3), none of the examined morphological or functional DTT trends were statistically different from Brownian motion (table S6). Further, trait dispersion of extant species around their MRCA was mostly continuous (fig. 4) with minimal evidence of clustering (i.e., discrete ecomorphs), a pattern largely consistent across radiations. One exception was found for *motion components* in Lake Victoria, which displayed a small secondary cluster of species with low values of jaw protrusion (fig. 4B), possibly representing divergence toward an adaptive peak associated with a substrate biting mode of feeding. That withstanding, comparably low trait variance in the two youngest radiations (fig. 2) are not suggestive of rapid divergence *among* lineages occupying distinct adaptive zones, where a significant portion of total potential diversity is achieved at initial stages of adaptive expansion.

Our study suggests that the diversification of feeding systems in cichlid adaptive radiations likely occurs by way of early burst, achieved not by adaptive divergence among clades, but through extremely rapid within-clade dispersion in incipient radiations. Previous research has predicted such patterns in cichlids as a possible outcome of transgressive segregation during widespread introgression – a common theme of emerging African lake radiations – paired with ecological opportunity in newly colonized habitats (Seehausen 2004; Meier et al. 2017; Irisarri 2018; Salzburger 2018; Meier et al. 2019; Selz and Seehausen 2019). Although we do not explicitly address the ecological dimensions across which functional diversification occurs, previous work on Lake Malawi and Tanganyika cichlids suggests that feeding diversity is distributed continuously along an axis of prey evasiveness (Martinez et al. 2018), matching observed patterns in this study of time-dependent trait dispersion around radiation specific MRCAs (fig. 4). If the landscape of ecological opportunity was discontinuously or sparsely distributed, for instance, it could pose challenges for a radiation diversifying via transgressive segregation since open adaptive zones are no longer adjacent to currently occupied zones, thereby reducing the probability that hybrid offspring happen upon a more distantly situated adaptive peak.

Clear evidence for early bursts appears to be the exception and not the rule in comparative trait data (Harmon et al. 2010). Previous analysis of trait diversification inclusive to Lake Tanganyika cichlids showed support for an early burst in body shape evolution but failed to find such evidence for jaw morphology (Ronco et al. 2021). Even this study, with rates among radiations showing a strong slowing trend with age, did not satisfy the prediction of an early burst by way of adaptive divergence among clades. Given that the latter is often used as a diagnostic tool for identifying early bursts (Harmon et al. 2003), it raises questions about potential limitations of our current models of adaptive radiation.

### Are we watching the same film?

Stephen Jay Gould famously contemplated what the diversity of life on earth might look like if we had the ability to start over and replay the tape of life (Gould 1991). Would it be unprecedented and unrecognizable, or would we see familiar patterns as selection inevitably leads to diversification along predictable paths? Many others have since pondered this question (e.g., Lobkovsky and Koonin 2012; Blount et al. 2018; Orgogozo 2015). In one sense, evolution is constantly repeating its own version of this experiment at much smaller spatial and temporal scales – independent radiations in related groups of organisms provide replication and insight into evolutionary contingencies under conditions of varying similarity. A primary focus of this study was to examine whether four large cichlid radiations, each resulting in hundreds of species and celebrated levels of ecological and morphological variation, have generated similar patterns of functional diversity. Has diversification of feeding functional morphology played out following the same script in each radiation, or have they diversified along separate functional axes? The answer appears to be that both are true.

Considering, for a moment, only the three African lake radiations examined in this study, there is an argument to be made that both functional and morphological diversification have progressed in a similar fashion in each of the lakes. The high-dimensional spaces filled by *motion components*, *motion pattern*, and cranial shape data each show the younger radiations, Malawi and Victoria, occupying subspaces of the older and more diverse Tanganyikan radiation, with novelty only commonplace in the latter. This result is consistent with impressions of widespread convergence on trophic morphotypes in Lakes Malawi and Tanganyika (Kocher et al. 1993; Ronco et al. 2021). On the other hand, it is unclear if we were to fast-forward the Lake Victoria tape to the current age of Lake Tanganyika (10-28 My, depending on the estimate), whether we would find a carbon-copy of that lake or if selection would eventually lead the Victorian radiation into unfamiliar functional and morphological spaces.

When we expand beyond the large African lake radiations to contrast their phenotypic and functional diversity for the first time with the large Neotropical radiation, a different story emerges. The two older radiations are more diverse than the others and each has invaded novel, radiation-specific regions of functional and morphological space. Tanganyika boasts species with unique combinations of *motion components*, including highly specialized planktivores (*Cyprichomis*) and benthic foragers (e.g., *Pseudosimochromis curvifrons*, *Telamatochromis vittatus*). In the Neotropics, some geophagine cichlids feed with a strongly sequenced kinematic pattern, partitioned between distinct jaw protrusion and cranial rotation phases (asymmetrical *motion patterns* toward the left of fig. 3B). Additionally, an innovation in select piscivorous species from the tribe Heroini (e.g., *Petenia splendida* and *Caquetaia myersi*) results in extreme levels of premaxillary protrusion (Waltzek and Wainwright 2003; Hulsey and García de León 2005) that places them in a unique region of *motion component* space (fig. 3A). Both Lake Tanganyika and the Neotropics contain cichlids with functional profiles that do not occur anywhere else, showing that radiations can eventually diverge from each other in key areas of diversification. These observations amplify questions about the contrasting landscapes of ecological opportunity experienced by lake versus continental radiations.

### Morphology provides an imperfect index of functional diversity

The idea that morphological variation can be used as a proxy for functional diversity is commonly advanced, but the widespread presence of complex form-function relationships tests this assumption (e.g., Wainwright et al. 2005; Young et al. 2007; 2010; Lautenschlager et al. 2020). Neotropical cichlids, when compared to the Lake Tanganyika radiation, illustrate that high variance in cranial morphologies does not always result in greater functional diversity. The primary axis of morphological variation in the Neotropics involved differences between elongate and slender (e.g., *Crenicichla*) versus deep heads with steep cranial profiles (e.g., *Symphysodon*, *Pterophyllum*, *Uaru*), which are adaptations typically found in species living in fast flowing riverine environments and slow-moving water like lakes or floodplains, respectively (López-Fernández et al. 2013). It is therefore likely that an important source of variance in head shapes of Neotropical cichlids is due to selection on habitat-specific body shape, perhaps involving adaptation of the locomotor system. Complex form-function relationships, particularly in biomechanical systems with many cooperating components, can make for challenging comparisons between morphologies and motions, and impact how these traits accumulate during adaptive diversification (Alfaro et al. 2005). These findings suggest caution is warranted in attributing observed morphological variation in fish feeding systems to functional diversity.

### Conclusions

Cichlids have captivated the attention of biologists and aquarists alike with their remarkable diversity, boasting seemingly endless combinations of body shapes, sizes, coloration patterns, diets, and behaviors. Each radiation examined in this study has amassed an impressive variety of morphological and functional diversity. Adaptive radiation of cichlids has produced modest diversity of feeding kinematics in Lakes Victoria and Malawi, while Lake Tanganyika has surpassed even the much older Neotropical radiation, suggesting that the forces driving diversification in Tanganyika outstrip those in the Neotropics. However, these patterns of diversity have been established on very different time scales. Rates of functional evolution range from 40-95 times faster in Lake Victoria than in the Neotropics, supporting the notion that the African Great Lake radiations have involved rapid evolutionary change. These observations suggest that while similarities exist, adaptive radiation of cichlid feeding kinematics has not always followed a common profile. Rather, evolutionary contingencies linked to time and biogeography explain varied patterns of morphological and functional diversification across this iconic group of fishes.

## Acknowledgements

We thank Ed Burress and Sarah Friedman for their support and assistance during this project. We also thank Michael Collyer for engaging discussions on morphometrics. CMM was funded by the UC Davis Chancellor’s Postdoctoral Fellowship Program. KAC was supported by an American Dissertation Fellowship from the American Association of University Women and a fellowship from the Achievement Rewards for College Scientists Foundation. ASRH was supported by the National Science Foundation with a Graduate Research Fellowship, Grant No. 1650042.

## Statement of Authorship

CMM and PCW conceived of the project. MDM and SRB recorded most of the videos of cichlid feeding events, and CMM contributed videos for the remaining species. All authors digitized landmark data on video frames. KAC ran statistical analyses. CMM and PCW wrote the paper with edits from all authors.

