## Supplemental Materials for "Replicated Functional Evolution in Cichlid Adaptive Radiations"

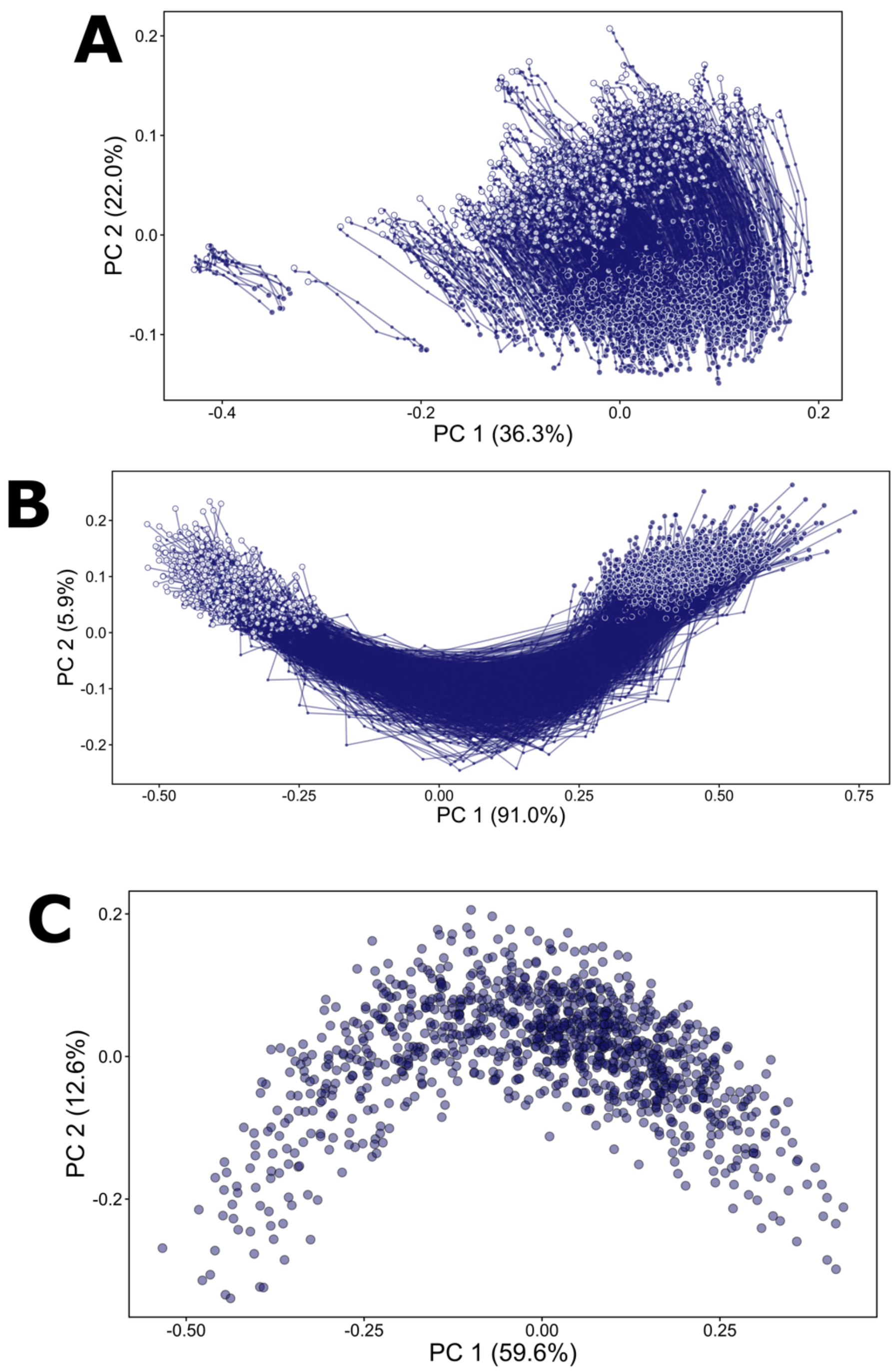


**Figure S1.** Process of alignment of shape trajectory data for the motion pattern trait. A) PCs 1 and 2 for aligned motion shapes representing 1,110 suction feeding events. Each line is a trajectory for a single feeding motion. The subjects of alignment (and scaling) are individual cranial shapes at different stages of a motion. B) Trajectory shapes shown in panel A are scaled and aligned. Here, the subject of alignment is the entire collection of 10 cranial shapes for a motion. C) Each of the 1,110 trajectory shapes is plotted as a single point in morphospace. Points closer to each other have more similar motion patterns than more distantly spaced points.

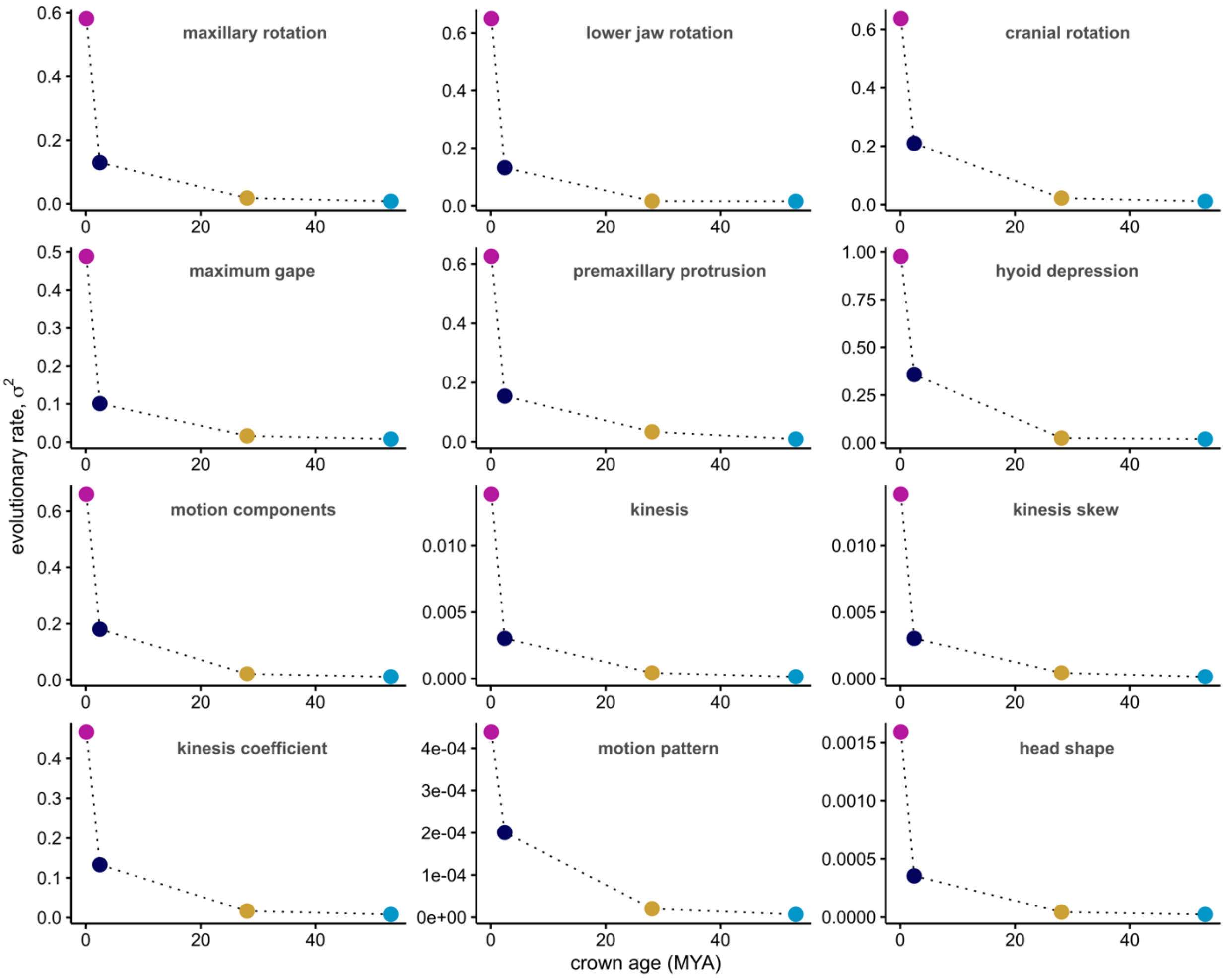


**Figure S2.** Rates of trait evolution versus radiation age. Estimated Brownian rate parameters for functional and morphological traits plotted against radiation crown age. Relative rates among radiations were highly consistent for all traits, all showing negative trends with radiation age.

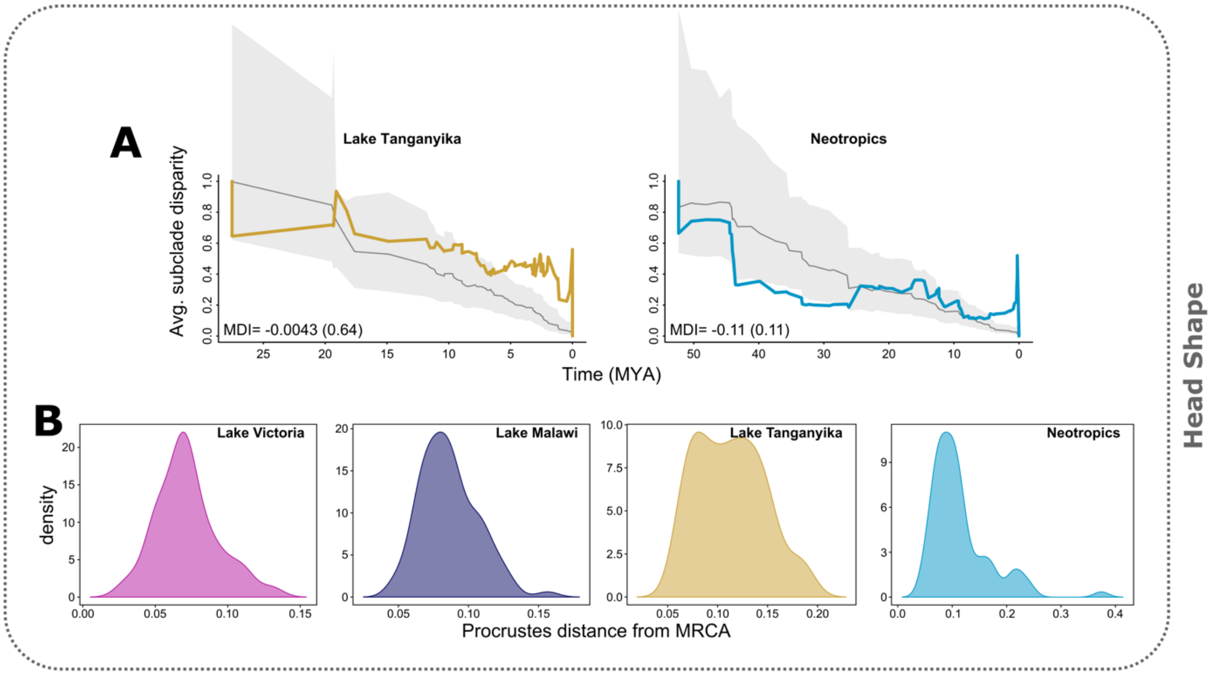


**Figure S3.** Evolution of head shape in cichlids. Disparity through time (DTT) plots for A) head shape in the two oldest cichlid radiations, Lake Tanganyika and the Neotropics. Also shown are B) distributions of Procrustes distances of head shapes between extant species and their most recent common ancestors (MRCA).

**Table S1.** Cichlid species examined in this study. 300 species are listed and colored by radiation. The number of individuals per species is provided in parentheses. On average, there were 3.1 (+/- 0.58 SD) feeding motions per individual fish. Asterisks denote species from Martinez et al. 2018. All other species represent new data to this study.

| | | |
|---|---|---|
| **LAKE VICTORIA** | Yssichromis sp. 'tipped blue' (1) | Lethrinops marginatus (1) |
| Enterochromis paropius (1) | **LAKE MALAWI** | Maylandia elegans (1) |
| Haplochromis chilotes (2) | Aristochromis christyi (2)* | Maylandia lombardoi (1) |
| Haplochromis howesi (1) | Aulonocara aquilonium (1) | Maylandia mbenjii (1) |
| Haplochromis lividus (1) | Aulonocara gertrudae (1) | Maylandia nigrodorsalis (1) |
| Haplochromis sp. 'blue obliquidens (1) | Aulonocara jacobfreibergi (1) | Mchenga conophoros (1) |
| Haplochromis sp. 'Murchison Bay' (1) | Aulonocara koningsi (1) | Melanochromis auratus (1)* |
| Haplochromis sp. 'thickskin' (1) | Aulonocara maylandi (1) | Melanochromis baliodigma (1) |
| Haplochromis thereuterion (1) | Aulonocara rostratum (1) | Melanochromis kaskazini (1)* |
| Haplochromis vonlinnei (1) | Aulonocara stuartgranti (1)* | Melanochromis loriae (1) |
| Harpagochromis cf. squamulatus (1) | Buccochromis nototaenia (1) | Melanochromis vermivorus (1) |
| Harpagochromis sp. 'golden duck' (1) | Buccochromis rhoadesii (1) | Melanochromis wochepa (1) |
| Labrochromis ishmaeli (1) | Buccochromis spectabilis (1) | Mylochromis anaphyrmus (1) |
| Lipochromis melanopterus (1) | Caprichromis liemi (1) | Mylochromis ericotaenia (1) |
| Lipochromis parvidens (1) | Champsochromis caeruleus (1) | Mylochromis sphaerodon (1) |
| Lipochromis sp. 'matumbi hunter' (1) | Champsochromis spilorhynchus (1) | Naevochromis chrysogaster (1) |
| Lithochromis rubripinnis (1) | Cheilochromis euchilus (1)* | Nimbochromis fuscotaeniatus (1) |
| Mbipia lutea (1) | Chilotilapia rhoadesii (1)* | Nimbochromis linni (1) |
| Neochromis greenwoodi (1) | Chindongo demasoni (1) | Nimbochromis polystigma (1)* |
| Neochromis nigricans (1) | Chindongo flavus (1)* | Nimbochromis venustus (1) |
| Neochromis omnicaeruleus (1) | Chindongo socolofi (1) | Nyassachromis boadzulu (1) |
| Neochromis rufocaudalis (1) | Copadichromis azureus (1) | Otopharynx antron (1) |
| Neochromis sp. 'Bihiru scraper' (1) | Copadichromis borleyi (3) | Otopharynx lithobates (1)* |
| Paralabidochromis chromogynos (1) | Copadichromis chrysonotus (3) | Otopharynx tetrastigma (1) |
| Paralabidochromis sauvagei (1) | Copadichromis geertsi (1) | Petrotilapia microgalana (1) |
| Platytaeniodus sp. 'red tail sheller' (1) | Copadichromis melas (1) | Placidochromis electra (1)* |
| Prognathochromis perrieri (1) | Copadichromis trewavasae (1) | Placidochromis johnstoni (1) |
| Ptyochromis fischeri (1) | Copadichromis virginalis (1) | Placidochromis milomo (2)* |
| Ptyochromis sp. 'red rock sheller' (1) | Ctenopharynx nitidus (1) | Placidochromis phenochilus (1) |
| Ptyochromis sp. 'salmon' (1) | Cynotilapia axelrodi (1) | Protomelas dejunctus (1) |
| Pundamilia macrocephala (1) | Cynotilapia zebroides (1) | Protomelas fenestratus (1) |
| Pundamilia nyererei (1) | Cyrtocara moorii (1) | Protomelas labridens (1) |
| Pundamilia pundamilia (1) | Dimidiochromis compressiceps (1) | Protomelas marginatus (1) |
| Pundamilia sp. 'crimson tide' (1) | Dimidiochromis kiwinge (1) | Protomelas spilonotus (1) |
| Pundamilia sp. 'pink anal' (1) | Fossorochromis rostratus (1)* | Protomelas spilopterus (1) |
| Pundamilia sp. 'red flank' (1) | Iodotropheus sprengerae (1) | Protomelas taeniolatus (1) |
| Pundamilia sp. 'red head' (1) | Labeotropheus fuelleborni (2)* | Pseudotropheus crabro (1)* |
| Pyxichromis orthostoma (1) | Labeotropheus trewavasae (2)* | Pseudotropheus cyaneorhabdos (1) |
| Xystichromis phytophagus (1) | Labidochromis caeruleus (2) | Pseudotropheus galanos (1) |
| Yssichromis piceatus (1) | Labidochromis joanjohsonae (1) | Pseudotropheus interruptus (1) |

**Table S1.** List of cichlid species, continued.

| | | |
|---|---|---|
| *Pseudotropheus johannii (1)* | *Lamprologus speciosus (1)* | *Trematochromis benthicola (2)** |
| *Rhamphochromis ferox (1)* | *Lepidiolamprologus attenuatus (3)** | *Triglachromis otostigma (1)* |
| *Rhamphochromis longiceps (1)** | *Lepidiolamprologus elongatus (1)* | *Tropheus brichardi (1)* |
| *Sciaenochromis fryeri (1)* | *Lepidiolamprologus nkambae (1)* | *Tropheus duboisi (1)* |
| *Stigmatochromis modestus (1)* | *Limnochromis auritus (2)** | *Tropheus moorii (1)* |
| *Taeniolethrinops laticeps (1)* | *Limnotilapia dardennii (1)* | *Variabilichromis moorii (2)** |
| *Tropheops romandi (1)* | *Lobochilotes labiatus (1)* | *Xenotilapia leptura (1)* |
| *Tyrannochromis nigriventer (1)* | *Neolamprologus boulengeri (2)** | *Xenotilapia melanogenys (1)* |
| **LAKE TANGANYIKA** | *Neolamprologus cylindricus (1)** | *Xenotilapia ochrogenys (1)* |
| *Altolamprologus calvus (1)* | *Neolamprologus fasciatus (1)** | *Xenotilapia ornatipinnis (1)* |
| *Altolamprologus compressiceps (1)* | *Neolamprologus furcifer (2)** | *Xenotilapia papilio (1)* |
| *Astatotilapia burtoni (1)** | *Neolamprologus hecqui (1)** | *Xenotilapia rotundiventralis (1)* |
| *Astatotilapia stappersii (1)* | *Neolamprologus longicaudatus (3)** | **NEOTROPICS** |
| *Aulonocranus dewindti (1)* | *Neolamprologus longior (3)** | *Acarichthys heckelii (1)* |
| *Bathybates minor (1)** | *Neolamprologus meeli (3)** | *Acaronia nassa (1)* |
| *Boulengerochromis microlepis (3)* | *Neolamprologus modestus (1)** | *Aequidens diadema (1)* |
| *Callochromis macrops (1)* | *Neolamprologus mondabu (1)* | *Aequidens metae (2)* |
| *Callochromis pleurospilus (1)* | *Neolamprologus multifasciatus (1)* | *Aequidens tetramerus (1)* |
| *Cardiopharynx schoutedeni (1)* | *Neolamprologus nigriventris (1)** | *Amatitlania kanna (1)* |
| *Chalinochromis brichardi (1)** | *Neolamprologus obscurus (4)** | *Amatitlania myrnae (1)* |
| *Chalinochromis popelini (1)* | *Neolamprologus prochilus (4)** | *Amatitlania nanolutea (1)* |
| *Ctenochromis horei (1)* | *Neolamprologus pulcher (1)** | *Amatitlania nigrofasciata (1)* |
| *Cyphotilapia frontosa (1)* | *Neolamprologus savoryi (3)** | *Amatitlania sajica (2)* |
| *Cyprichromis coloratus (1)* | *Neolamprologus sexfasciatus (2)** | *Amatitlania septemfasciata (1)* |
| *Cyprichromis leptosoma (1)* | *Neolamprologus similis (1)* | *Amatitlania siquia (1)* |
| *Cyprichromis microlepidotus (1)* | *Neolamprologus tetracanthus (1)** | *Amphilophus citrinellus (1)* |
| *Cyprichromis pavo (1)** | *Neolamprologus walteri (1)** | *Amphilophus labiatus (1)* |
| *Cyprichromis zonatus (1)* | *Ophthalmotilapia boops (1)** | *Amphilophus sagittae (1)* |
| *Ectodus descampsii (1)** | *Ophthalmotilapia nasuta (3)** | *Amphilophus trimaculatus (1)* |
| *Eretmodus cyanostictus (1)** | *Paracyprichromis brieni (1)* | *Andinoacara stalsbergi (1)* |
| *Gnathochromis permaxillaris (1)** | *Paracyprichromis nigripinnis (1)* | *Archocentrus centrarchus (1)* |
| *Gnathochromis pfefferi (1)* | *Perissodus microlepis (2)* | *Astronotus ocellatus (1)* |
| *Grammatotria lemairii (1)* | *Petrochromis orthognathus (1)* | *Australoheros scitulus (1)* |
| *Greenwoodochromis bellcrossi (1)* | *Pseudosimochromis babaulti (1)** | *Biotodoma cupido (1)* |
| *Haplotaxodon microlepis (1)** | *Pseudosimochromis curvifrons (1)* | *Biotodoma wavrini (1)* |
| *Interochromis loocki (1)* | *Reganochromis calliurus (1)* | *Caquetaia kraussii (1)* |
| *Julidochromis dickfeldi (1)** | *Spathodus erythrodon (1)* | *Caquetaia myersi (3)* |
| *Julidochromis marlieri (1)* | *Tanganicodus irsacae (1)* | *Caquetaia spectabilis (1)* |
| *Julidochromis ornatus (1)* | *Telmatochromis brichardi (1)* | *Chaetobranchus flavescens (1)* |
| *Lamprologus callipterus (1)** | *Telmatochromis dhonti (1)** | *Chiapaheros grammodes (1)* |
| *Lamprologus lemairii (3)** | *Telmatochromis temporalis (2)** | *Cichla ocellaris (1)* |
| *Lamprologus ornatipinnis (1)* | *Telmatochromis vittatus (1)* | *Cichlasoma dimerus (1)* |
| *Lamprologus signatus (1)* | *Trematocara variabile (3)* | *Cincelichthys bocourti (1)* |

**Table S1.** List of cichlid species, continued.

| | | |
|---|---|---|
| *Cleithracara maronii (1)* | *Heroina isonycterina (1)* | *Parachromis dovii (1)* |
| *Crenicara punctulatum (1)* | *Heros severus (2)* | *Parachromis managuensis (1)* |
| *Crenicichla lepidota (1)* | *Herotilapia multispinosa (1)* | *Paraneetroplus gibbiceps (1)* |
| *Crenicichla regani (1)* | *Hoplarchus psittacus (1)* | *Petenia splendida (1)* |
| *Crenicichla saxatilis (1)* | *Hypselecara coryphaenoides (1)* | *Pterophyllum scalare (3)* |
| *Crenicichla strigata (1)* | *Hypselecara temporalis (1)* | *Retroculus lapidifer (1)* |
| *Crenicichla sveni (1)* | *Hypsophrys nematopus (1)* | *Rocio octofasciata (1)* |
| *Cribroheros robertsoni (1)* | *Krobia xinguensis (1)* | *Satanoperca daemon (1)* |
| *Cryptoheros spilurus (1)* | *Kronoheros umbrifer (1)* | *Symphysodon discus (3)* |
| *Dicrossus maculatus (1)* | *Laetacara araguaiae (1)* | *Tahuantinsuyoa macantzatza (1)* |
| *Geophagus abalios (1)* | *Laetacara thayeri (1)* | *Talamancaheros sieboldii (1)* |
| *Geophagus crassilabris (1)* | *Maskaheros argenteus (1)* | *Teleocichla centrarchus (1)* |
| *Geophagus iporangensis (1)* | *Mayaheros urophthalmus (3)* | *Trichromis salvini (3)* |
| *Geophagus megasema (1)* | *Mesoheros festae (1)* | *Uaru amphiacanthoides (1)* |
| *Geophagus parnaibae (1)* | *Mesonauta mirificus (1)* | *Vieja breidohri (1)* |
| *Geophagus steindachneri (1)* | *Mikrogeophagus ramirezi (1)* | *Vieja fenestrata (2)* |
| *Geophagus winemilleri (1)* | *Nandopsis tetracanthus (2)* | *Vieja synspila (1)* |
| *Guianacara stergiosi (1)* | *Nosferatu pantostictus (1)* | |
| *Gymnogeophagus tiraparae (1)* | *Panamius panamensis (1)* | |

**Table S2**. Phylogenetic placement of undescribed Lake Victoria cichlids. Correspondence of undescribed species from this study with the tree tips used from a published phylogeny (McGee et al. 2020). Sources of supporting literature for placements are also provided. See supplemental methods for additional procedural details.

| This Study | McGee et al. (2020) Phylogeny | Source |
|---|---|---|
| *Haplochromis sp. 'blue obliquidens'* | *Haplochromis obliquidens* | Seehausen (1996) |
| *Haplochromis sp. 'Murchison Bay'* | *Haplochromis riponianus* | Greenwood (1980) |
| *Haplochromis sp 'thickskin'* | *Haplochromis plagiodon* | McGee et al. (2020) |
| *Harpagochromis sp. 'golden duck'* | *Haplochromis prognathus* | Greenwood (1980) |
| *Lipochromis sp. 'matumbi hunter'* | *Haplochromis obesus* | Seehausen (1996) |
| *Neochromis sp. 'Bihiru scraper'* | *Neochromis simotes* | Seehausen (1996) |
| *Platytaeniodus sp. 'red tail sheller'* | *Haplochromis xenognathus* | Greenwood (1980) |
| *Ptyochromis sp. 'red rock sheller'* | *Haplochromis granti* | Greenwood (1980) |
| *Ptyochromis sp. 'salmon'* | *Haplochromis prodromus* | Greenwood (1980) |
| *Pundamilia sp. 'crimson tide'* | *Pundamilia igneopinnis* | Seehausen et al. (1998) |
| *Pundamilia sp. 'pink anal'* | *Mbipia mbipi* | McGee et al. (2020) |
| *Pundamilia sp. 'red flank'* | *Haplochromis parorthostoma* | Seehausen (1996) |
| *Pundamilia sp. 'red head'* | *Haplochromis pallidus* | Seehausen (1996) |
| *Yssichromis sp. 'tipped blue'* | *Haplochromis pyrrhocephalus* | Greenwood (1980) |

**TABLE S3.** Comparisons of variances for functional traits and head shape. Variances are shown on the left and P-values from pairwise comparisons are on the right. Statisically significant results are shown in bold, and are based on 10,000 permutations

| | **Variances** | | | | **Pairwise P-values** | | | | | |
|---|---|---|---|---|---|---|---|---|---|---|
| **Trait** | **Vic** | **Mal** | **Tan** | **Neo** | **Mal - Neo** | **Mal - Tan** | **Mal - Vic** | **Neo - Tan** | **Neo - Vic** | **Tan - Vic** |
| motion components | 0.38 | 0.60 | 1.22 | 1.10 | **0.00040** | **0.00010** | 0.21 | 0.43 | **0.00010** | **0.00010** |
| motion pattern | 0.014 | 0.031 | 0.042 | 0.033 | 0.82 | 0.064 | **0.016** | 0.10 | **0.0094** | **0.00040** |
| premaxillary protrusion | 0.058 | 0.11 | 0.27 | 0.17 | 0.14 | **0.00010** | 0.30 | **0.017** | **0.029** | **0.00020** |
| maxillary rotation | 0.056 | 0.066 | 0.15 | 0.090 | 0.34 | **0.00050** | 0.74 | **0.012** | 0.27 | **0.0022** |
| lower jaw rotation | 0.065 | 0.074 | 0.17 | 0.15 | **0.0070** | **0.00070** | 0.82 | 0.43 | **0.019** | **0.0030** |
| cranial rotation | 0.058 | 0.10 | 0.20 | 0.27 | **0.00020** | **0.032** | 0.46 | 0.15 | **0.00060** | **0.017** |
| hyoid depression | 0.091 | 0.20 | 0.29 | 0.34 | **0.0054** | 0.091 | 0.098 | 0.26 | **0.00030** | **0.0034** |
| mouth gape | 0.047 | 0.051 | 0.14 | 0.084 | 0.33 | **0.0020** | 0.91 | 0.080 | 0.35 | **0.022** |
| kinesis | 0.0014 | 0.0016 | 0.0040 | 0.0021 | 0.44 | **0.00010** | 0.70 | **0.0007** | 0.31 | **0.00080** |
| log-kinesis skew | 0.011 | 0.030 | 0.047 | 0.036 | 0.49 | **0.034** | 0.055 | 0.16 | **0.011** | **0.00050** |
| log-kinesis coefficient | 0.043 | 0.061 | 0.12 | 0.17 | **0.0003** | 0.055 | 0.65 | 0.15 | **0.0022** | **0.046** |
| head shape | 0.0055 | 0.0066 | 0.013 | 0.015 | **0.00010** | **0.00060** | 0.63 | 0.25 | **0.00020** | **0.0022** |

Vic= Lake Victoria; Mal= Lake Malawi; Tan= Lake Tanganyika; Neo=Neotropics

**Table S4.** Hypervolume results for functional and morphological traits. Each row contains a comparison between two radiations or between one radiation and all other species. The column to the right displays results of a randomization procedure comparing fraction of unique hypervolume spaces bewteen cichlid radiations. Percentiles display how extreme observed results are relative to 10,000 randomized permutations.

| Trait | Hypervolume 1 (H1) | Hypervolume 2 (H2) | Volume (H1, H2) | Sorensen overlap | Unique fraction (H1, H2) | Percentile from Randomization: Unique fraction (H1, H2) |
|---|---|---|---|---|---|---|
| components | Tanganyika | Non-Tanganyika | (14.93, 7.00) | 0.52 | (0.62, 0.19) | (0.999, 0.096) |
| components | Neotropics | Non-Neotropics | (9.50, 7.87) | 0.56 | (0.49, 0.39) | (0.995, 0.797) |
| components | Malawi | Non-Malawi | (3.74, 12.51) | 0.43 | (0.06, 0.72) | (0.000, 0.999) |
| components | Victoria | Non-Victoria | (1.77, 11.34) | 0.26 | (0.03, 0.85) | (0.000, 0.999) |
| components | Tanganyika | Neotropics | (14.93, 9.50) | 0.51 | (0.58, 0.35) | (0.996, 0.695) |
| components | Tanganyika | Malawi | (14.93, 3.74) | 0.37 | (0.77, 0.08) | (0.999, 0.001) |
| components | Tanganyika | Victoria | (14.93, 1.77) | 0.21 | (0.88, 0.01) | (0.999, 0.999) |
| components | Malawi | Victoria | (3.74, 1.77) | 0.53 | (0.61, 0.17) | (0.971, 0.117) |
| components | Malawi | Neotropics | (3.74, 9.50) | 0.44 | (0.23, 0.69) | (0.232, 0.999) |
| components | Victoria | Neotropics | (1.77, 9.50) | 0.25 | (0.22, 0.85) | (0.257, 0.999) |
| motion pattern | Tanganyika | Non-Tanganyika | (0.0078, 0.0038) | 0.62 | (0.54, 0.050) | (0.997, 0.001) |
| motion pattern | Neotropics | Non-Neotropics | (0.0047, 0.0047) | 0.74 | (0.26, 0.26) | (0.632, 0.595) |
| motion pattern | Malawi | Non-Malawi | (0.0042, 0.0052) | 0.78 | (0.14, 0.30) | (0.151, 0.707) |
| motion pattern | Victoria | Non-Victoria | (0.0013, 0.0055) | 0.38 | (0.010, 0.76) | (0.999, 0.999) |
| motion pattern | Tanganyika | Neotropics | (0.0078, 0.0047) | 0.66 | (0.47, 0.12) | (0.941, 0.080) |
| motion pattern | Tanganyika | Malawi | (0.0078, 0.0042) | 0.66 | (0.49, 0.060) | (0.960, 0.004) |
| motion pattern | Tanganyika | Victoria | (0.0078, 0.0013) | 0.29 | (0.83, 0.010) | (0.999, 0.999) |
| motion pattern | Malawi | Victoria | (0.0042, 0.0013) | 0.47 | (0.69, 0.030) | (0.996, 0.001) |
| motion pattern | Malawi | Neotropics | (0.0042, 0.0047) | 0.73 | (0.22, 0.31) | (0.422, 0.659) |
| motion pattern | Victoria | Neotropics | (0.0013, 0.0047) | 0.42 | (0.040, 0.73) | (0.004, 0.998) |
| head shape | Tanganyika | Non-Tanganyika | (0.0014, 0.00088) | 0.53 | (0.57, 0.3) | (0.999, 0.672) |
| head shape | Neotropics | Non-Neotropics | (0.0011, 0.00083) | 0.46 | (0.59, 0.47) | (0.999, 0.982) |
| head shape | Malawi | Non-Malawi | (0.00033, 0.0016) | 0.33 | (0.05, 0.8) | (0.000, 0.999) |
| head shape | Victoria | Non-Victoria | (0.00022, 0.0014) | 0.27 | (0.03, 0.84) | (0.000, 0.999) |
| head shape | Tanganyika | Neotropics | (0.0014, 0.0011) | 0.44 | (0.62, 0.48) | (0.999, 0.976) |
| head shape | Tanganyika | Malawi | (0.0014, 0.00033) | 0.33 | (0.8, 0.13) | (0.999, 0.041) |
| head shape | Tanganyika | Victoria | (0.0014, 0.00022) | 0.25 | (0.86, 0.08) | (0.999, 0.013) |
| head shape | Malawi | Victoria | (0.00033, 0.00022) | 0.62 | (0.49, 0.23) | (0.898, 0.385) |
| head shape | Malawi | Neotropics | (0.00033, 0.0011) | 0.33 | (0.3, 0.78) | (0.648, 0.999) |
| head shape | Victoria | Neotropics | (0.00022, 0.0011) | 0.26 | (0.26, 0.84) | (0.483, 0.999) |

**TABLE S5.** Comparisons of **e**volutionary rates for functional traits and head shape. Rates are shown on the left and P-values from pairwise comparisons are on the right. Statisically significant results are shown in bold, and are based on 10,000 permutations.

| | **Brownian rate parameter** | | | | **Pairwise P-values** | | | | | |
|---|---|---|---|---|---|---|---|---|---|---|
| **Trait** | **Vic** | **Mal** | **Tan** | **Neo** | **Mal - Neo** | **Mal - Tan** | **Mal - Vic** | **Neo - Tan** | **Neo - Vic** | **Tan - Vic** |
| motion components | 0.66 | 0.18 | 0.022 | 0.012 | **0.00020** | **0.00015** | **0.0012** | **0.046** | **0.00010** | **0.00010** |
| motion pattern | 0.00044 | 0.00020 | 0.000020 | 0.0000068 | **0.00020** | **0.00010** | **0.040** | **0.00040** | **0.00010** | **0.00010** |
| premaxillary protrusion | 0.63 | 0.15 | 0.033 | 0.0091 | **0.00020** | **0.00090** | **0.021** | **0.0060** | **0.00020** | **0.00020** |
| maxillary rotation | 0.58 | 0.13 | 0.018 | 0.0079 | **0.00020** | **0.00020** | **0.010** | 0.076 | **0.00020** | **0.00015** |
| lower jaw rotation | 0.65 | 0.13 | 0.016 | 0.015 | **0.00020** | **0.00015** | **0.0084** | 0.96 | **0.00020** | **0.00010** |
| cranial rotation | 0.64 | 0.21 | 0.022 | 0.011 | **0.00020** | **0.00020** | **0.040** | 0.13 | **0.00010** | **0.00010** |
| hyoid depression | 0.98 | 0.36 | 0.025 | 0.020 | **0.00020** | **0.00020** | 0.054 | 0.57 | **0.00010** | **0.00010** |
| mouth gape | 0.49 | 0.10 | 0.016 | 0.0078 | **0.00010** | **0.00010** | **0.0057** | 0.12 | **0.00010** | **0.00010** |
| kinesis | 0.014 | 0.0030 | 0.00043 | 0.00015 | **0.00020** | **0.00010** | **0.0062** | **0.017** | **0.00015** | **0.00010** |
| log-kinesis skew | 0.12 | 0.070 | 0.010 | 0.0025 | **0.00010** | **0.00010** | 0.21 | **0.00015** | **0.00020** | **0.00020** |
| log-kinesis coefficient | 0.47 | 0.13 | 0.017 | 0.0080 | **0.00020** | **0.00015** | **0.027** | 0.11 | **0.00020** | **0.00020** |
| head shape | 0.0016 | 0.00035 | 0.000043 | 0.000023 | **0.00020** | **0.00015** | **0.00020** | **0.03** | **0.00010** | **0.00010** |

Vic= Lake Victoria; Mal= Lake Malawi; Tan= Lake Tanganyika; Neo=Neotropics

**Table S6.** Results from Disparity Though Time (DTT) analyses. Morphological Disparity Inde (MDI) values are reported, describing the deviation of subclade trait disparities from simulated values under Brownian motion. P-values shown in parentheses. Note that none of the results are statistically significant, meaning that trait diversification was statistically consistent with a Brownian motion model of evolution.

| Trait | All Species | Victoria | Malawi | Tanganyika | Neotropics |
|---|---|---|---|---|---|
| motion components | 0.029 (0.72) | -0.012 (0.37) | 0.15 (0.92) | 0.0088 (0.61) | 0.019 (0.69) |
| motion pattern | 0.096 (0.88) | 0.12 (0.84) | 0.40 (0.99) | 0.30 (0.98) | 0.16 (0.94) |
| head shape | -0.083 (0.40) | -0.080 (0.067) | 0.092 (0.84) | -0.0043 (0.64) | -0.11 (0.11) |

### Supplementary Information

#### Materials & Methods

*Phylogeny matching* – For comparative analyses, species were matched to a phylogeny of cichlids by McGee and colleagues (McGee et al. 2020). This phylogeny is limited to only described cichlid species. Several cichlids from Lake Victoria in our dataset (n=14 species) are currently known in the literature but remain undescribed (e.g., Pundamilia sp. 'red flank'). We used existing phylogenomic and systematic knowledge to assign undescribed taxa to Lake Victoria cichlid species in the phylogeny with which they share key diagnostic features or previous molecular studies have suggested may be close relatives (Table S2; Greenwood 1980; Seehausen 1996; Seehausen et al. 1998; McGee et al. 2020).

*Crown ages of cichlid radiations* – There is considerable interest in the age of cichlids and of major clades within this group. Possible ages of the two youngest radiations in this study, Lakes Malawi and Victoria, have traditionally been bounded by geological events. Lake Malawi is reported to have almost completely desiccated between 1.6-1 Ma (Delvaux 1995). Additionally, Lake Victoria cichlids are thought to have radiated within the last 150,000 years (Meier et al. 2017), although evidence exists for large-scale (and possibly complete) desiccation of the lake as recently as 18,000-10,000 years ago (Stager and Johnson 2008). For the Lake Tanganyika radiation, estimates on the order of 30-20 Ma (e.g., McGee et al. 2020) have recently been challenged by a phylogenomic assessment placing the age of the radiation at around 10 Ma (Ronco et al. 2021). Additionally, there has been much debate and uncertainty surrounding the timing of the split between the subfamily Cichlinae (Neotropical cichlids) and Pseudocrenilabrinae (African cichlids), and by extension, the crown ages of these large clades.

Some earlier molecular-based phylogenetic reconstructions resulted in topologies that appeared congruent with the splitting of landmasses during Gondwanan vicariance, which would place divergence of current day Neotropical and African subfamilies at 120-100 Ma (Sparks 2004; Sparks and Smith 2004). However, several recent studies favor a trans-Atlantic dispersal model of cichlid phylogeography, with several estimates between 70-45 Ma (Matschiner 2018). Moreover, an even more recent study using whole genomes and fossil calibration places the divergence of Cichlinae and Pseudocrenilabrinae at approximately 62 Ma (Matschiner et al. 2020).

The phylogeny used in this study was trimmed from a previously published tree (McGee et al. 2020), although displaying some differences from ages listed above, preserves a rank order of radiation ages that is consistent with recent phylogenetic reconstructions of cichlids; with an estimated 53.1 Ma for the Neotropics, 28.1 Ma for Lake Tanganyika, 2.4 Ma for Lake Malawi, and 0.1 Ma for Lake Victoria. While the accepted timing of cichlid evolution can (and certainly will) change with new data, short of a major upheaval in the estimates of divergence times, this order of crown ages is unlikely to be upturned. Therefore, rates of trait evolution examined in our study, which depend on inferred timing from a phylogeny, may similarly change, but overall interpretations of functional diversification across radiations differing in their relative time scales (tens of millions of years to thousands of years) are expected to remain largely intact.
